# Searching for mates may shape the immune response and parental investments

**DOI:** 10.1101/2022.05.08.491084

**Authors:** Hasan Sevgili

## Abstract

In the sexual reproduction model, a wide variety of strategies have emerged in different animal species for individuals to pursue the opposite sex for mating purposes. Many ways (such as chemical signals and calling songs) are utilized to stimulate or invite the other sex. This may lead to differential evolution of sexes in a species. For example, the limitations in signal production or perception of one of the sexes can make this sex sedentary, therefore strategies in pursuing mates evolve diversely among sexes. In this context, McCartney et al. (2012) proposed a theory to explain the differentiation of the mate-searching system in some species of *Poecilimon* bushcrickets and the reasons for the change in the sexual roles in the search. The aim of the present study was the testing of this theory proposed by McCartney et al. (2012) for *Poecilimon* species on spermatophore investment, with two species (*P. sureyanus*-female searching and *P. inflatus*- male searching), which have different strategies in terms of females’ response to male calling songs. Reproductive effort and physiological activities are both costly for the organism, and these traits belie a trade-off due to the limited energetic resources. Therefore, some immune parameters (Phenoloxidase and lytic activities) of these two species were compared within the context of the allocation of resources resulting from the trade-off between reproductive functions and immune defence. Immune activities are much more plastic in nature than the differentiation of reproductive traits among sexes. The results of this study showed that a mate searching strategy difference, based on the female’s ability to respond to male call sounds -which must acoustically communicate for reproduction-may play a decisive role in both spermatophore and immune activities. On the other hand, the findings also indicated that there may be trade-offs between the costs related to reproduction and the immune system. In addition, it has been revealed that there may be a trade-off between the immune activities and the differences between sexes. On the other hand, our data indicated that the duration of morphogenesis did not have a decisive role on spermatophore and sperm allocation in bushcrickets.

## Introduction

The energy costs of reproduction, mate searching and survival during the mating period in insects have led to the evolution of different strategies between species according to the role of the sex. The methods used to find a potential mate (calling songs, pheromone production, advertisements, displays, mobility) show a significant dimorphism between sexes, and this difference naturally places the cost of reproductive investment more on one of the sexes. In general, a significant portion of the burden of the pre-reproductive activity is put on the male’s tab (Simmons 2001; Fromhage et al. 2016).

In a population, mating behaviors between sexes (e.g. alternative mating tactics) have evolved along with stable strategies (Cowden 2012) which emerged in the evolutionary process (sexual selection). While pre-mating, mate searching, and evolved strategies related to this process stabilise the behaviour of sexes in populations of the same species, these strategies may differ in this context between close relatives of species (Thornhill 1979; McCartney et al. 2012).

The answers to the question of how asymmetrical strategies in parental investment regarding reproduction may differ in species within the same genus as different evolutionarily stable strategies, have come from the brachypterous genus *Poecilimon* (bushcrickets, Orthoptera) which exhibits high species richness (McCartney et al. 2012). The species of the genus have served as good models in evolutionary biology.

In bush-crickets such as the genera *Isophya* and *Poecilimon* (Orthoptera: Tettigoniidae), males transfer both a sperm-containing ampulla and the spermatophylax surrounding it (ampulla +spermatophylax= Spermatophore) to the female during mating (Mccartney et al. 2008; Lehmann 2012; Uma and Sevgili 2014). Spermatophore weight and content (sperm count) vary widely between (or even within) species (Mccartney et al. 2008; Sevgili et al. 2015).

At the same time, there exist different strategies (such as mating multiple times with different males) to obtain spermatophore, which is a very valuable source of sustenance for the females. The quality of females is an important indicator in mate searching males, and they generally tend to invest more sperm and spermatophore to larger females (Uma and Sevgili 2014). In living organisms, various strategies have evolved to allow the allocation of energy, which is a limited resource, to reproductive (egg laying and offspring care) and pre-reproductive efforts (calling song, nuptial gifts), alongside necessary activities of survival (such as immunity).

In addition to the costly spermatophore production, male’s effort to produce the necessary acoustic signals requires a significant amount of energy. The male calling song has important indicator signals of the male’s body size which is positively correlated with a larger spermatophore, which provides more nutrition and better potential mate selection (Lehmann and Lehmann 2008). *Poecilimon* bush-crickets have two different communication systems. In some species, mute females approach the males phonotactically, while in the others females respond acoustically to calling songs of males and thus can actively signal the male to approach them phonotactically (Heller and von-Helversen 1993). The differentiation of the mate searching strategy within the same genus has led to the evolution of a trade-off in the spermatophore investment of the male within these two communication systems (McCartney et al. 2012).

One of the most important effects of the active sex role in mate searching is the distribution of the cost of mating (sound producing, male’s produce the nuptial gifts, etc.) according to benefit and cost. In the system where the males search for the signal-producing female, generally smaller spermatophores are produced than in the other mate-searching system (McCartney et al. 2012). The cost of the potential-mate-searching role that generates the signals brings with it survival risks, by adjusting spermatophore size, and also by becoming more susceptible to predation. Predation risks of the sex which reveals its own position by producing a sound signal (usually males) and the mute sex which is mobilized to locate the signal source may also be different (Heller 1992a; Raghuram et al. 2015; but see Torsekar et al. 2019).

While evolutionarily stable strategies that differ between species shape behaviours dedicated to the reproductive costs of mating, questions about how physiological mechanisms for survival may be affected in this process have not been adequately answered. It has been reported that the immune system, which is one of the most important physiological characteristics related to survival, is significantly affected by the cost of mating and reproductive behaviours that can differ between species (Fedorka et al. 2004; Nunn et al. 2009; Kerr et al. 2010).

Many studies have reported that males and females show dimorphism in the context of immune investment, and in general, females have stronger immunity in insects (Rolff 2002; Nunn et al. 2009; Kelly et al. 2018). However, this general acceptance is not valid for all immune parameters, and a few studies have reported that some immune responses in males are more active than females (Fedorka et al. 2004; Kelly 2017; Sevgili 2019a; Kirschman et al. 2019).

This indicates that sexual dimorphism has a trade-off over physiological costs in terms of reproduction and other survival strategies. Immune competence and reproductive effort (such as call song and spermatophore production) are two important traits that require energy and show trade-off characteristics especially in the adult period (Rolff and Siva-Jothy 2002; Fedorka and Sevgili 2014).

If the reproductive cost of two different mate searching systems in *Poecilimon* species led to the emergence of different strategies in spermatophore weight and sperm count in these two groups (McCartney et al. 2012), whether immune responses are shaped according to the same strategy along with reproductive parameters due to physiological cost is an important question. Because studies on many insect species have shown that there is a trade-off between physiological traits such as reproductive investments, developmental process and immune competence related to survival (Fedorka et al. 2004; Kelly 2011; Mcnamara et al. 2013; Barbosa et al. 2016). All these trade-off processes try to balance both reproduction and survival within the population by improving the individual’s fitness.

The species of the genus *Poecilimon* generally have a life span of 2-3 months after hatching, and when they enter the very short adult period, they must mate and reproduce quickly besides their struggle for survival. In this study, the proposed theory of spermatophore investment (McCartney et al. 2012), and whether this theory can be supported by some immune activities were investigated on two *Poecilimon* species, which have different strategies in terms of females’ response to male calling song. Immune activities have a much more plastic characteristic than reproductive traits that differ significantly between sexes. On the other hand, while there is a relationship between increased reproductive success and suppression of immunity (Rolff and Siva-Jothy 2002), it is expected that immunity will be weaker in species that produce larger spermatophores and more sperm.

## Material and Methods

### The insects

In this study, two endemic species belonging to the genus *Poecilimon*, in which the mate searching system differs according to whether the females can respond vocally to male calling songs (McCartney et al. 2012), were utilised. One of them, *Poecilimon (Poecilimon) sureyanus* Uvarov, which is known from Northwest Anatolia, was collected from Uludağ University Campus (N 40.2327, E 28.8808, 16.06.2017, Bursa, TR). Since the tegmina are atrophied in females of this species, a sound signal cannot be generated (Fig. 1). The other species is *Poecilimon (Poecilimon) inflatus Brunner von Wattenwyl*, which is distributed in the Western Mediterranean and Southern Aegean regions, was collected as adults from around Dalyan (N 36.7686, E 28.6847, 14.05.2016, Muğla, TR). In this species, females respond to male calling songs (Fig. 1) as their wings are developed and are in contact with each other dorsally. Sample sound recordings of both species were made with Condenser Ultrasound Microphone (Avisoft-Bioacoustics CM16/CMPA-P48, sampling rate 192 KHz, 16 bit) connected to TASCAM HD-P2. Since these two species were collected as adults from their habitats, their mating histories were unknown.

**Figure 1.**
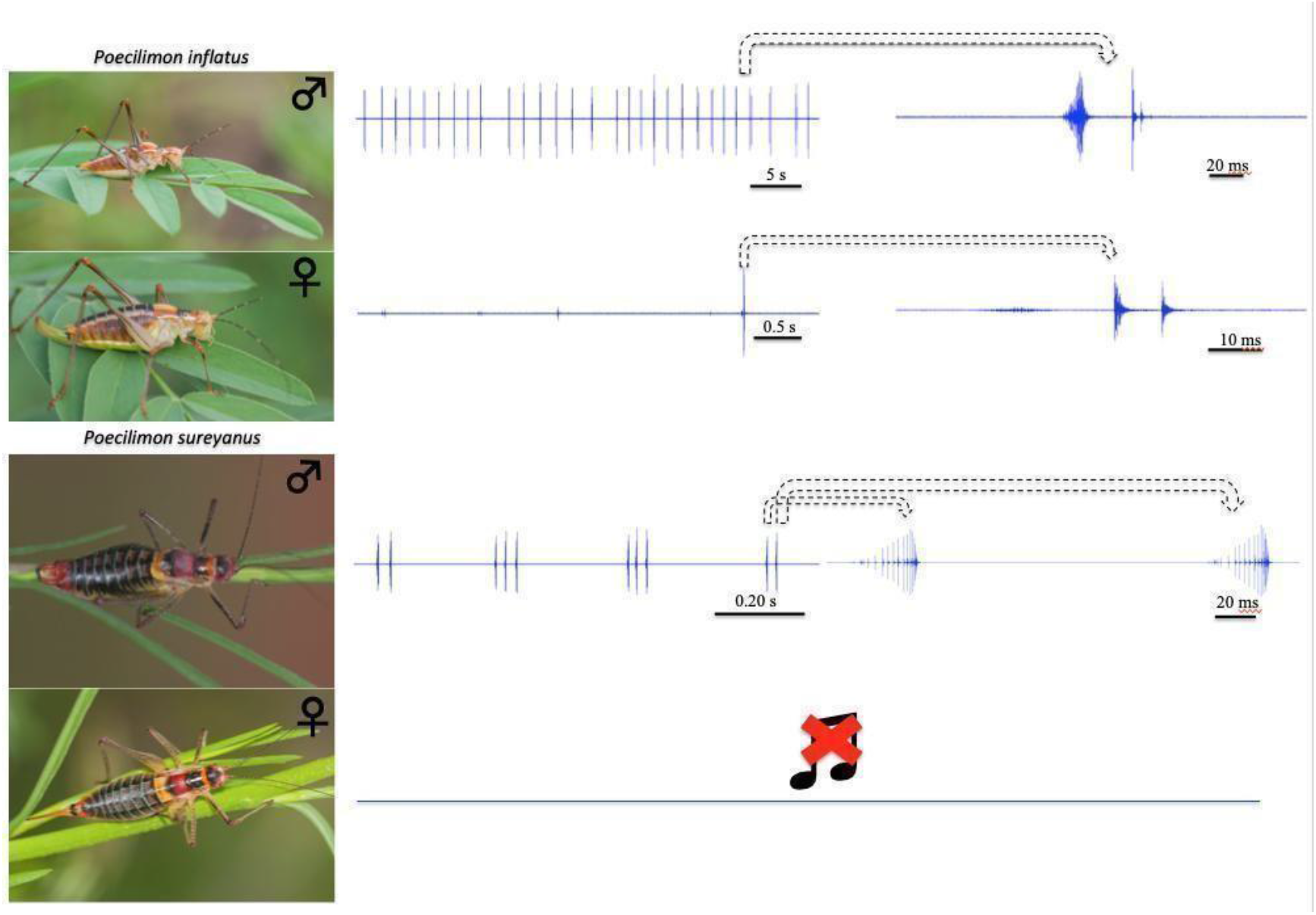
In *P. inflatus*, which can typically be distinguished from other similar groups by the swollen posterior part of the pronotum, females can respond to male calls. However, *P. sureyanus* females with a reduced wing structure that are not suitable for producing sound are mute.

To know the mating history of the bush-crickets, individuals were provided to mate with 5 males and 5 females in 20×20 (cm) cages in which fresh nettle, dandelion, apple and cucumber slices were placed under laboratory conditions (±25°C). Cages were cleaned at regular intervals and their food was replaced with fresh ones. Petri dishes containing sand mixed with soil were placed inside the cages to collect the eggs. These petri dishes containing eggs were kept in laboratory conditions throughout the summer and were sprayed with water from time to time to provide humidity. From autumn, these petri dishes were taken into climate chambers with controlled humidity (around 60-70%), and the temperature was reduced every 15 days in accordance with the seasonal change in the climate of the region, and wintering was induced at 6-7°C in the winter season. During this period, water was sprayed on the petri dishes once a week. After February, temperature was adjusted to 10°C, 15°C, and 18°C, respectively, every 10 days. Eggs started to hatch from the beginning of March and nymphs were collected daily with the help of a painting brush. 94 *P. sureyanus* and 93 *P. inflatus* nymphs were obtained from the eggs. However, 54 *P. sureyanus* and 47 *P. inflatus* reached the adult stage. Nymphs were initially reared in small plastic bottles (250 ml) with nettle and cucumber slices. Bottles and food were replaced every two days. Daily recordings were made from egg hatching to adulthood of individuals of each species (Development duration). The sexes (two species separately) were taken into separate clean cages to prevent mating of individuals that became last instar nymphs to be used in the experiment.

### Mating and immune assays

The methodology of bush-cricket mating trials given in the related literature (Uma and Sevgili 2014; Yiǧit et al. 2019) was followed for weighing individuals (before and after mating), copulation, determination of spermatophore characteristics (weighing) and sperm counting. All mated individuals were virgins. One week after mating, randomly selected males and females from cages were tagged. In previous studies, this one-week waiting period was determined as the ideal mating age after the male matured to produce enough sperm in Barbistini bush-crickets (Uma and Sevgili 2014; Sevgili et al. 2015; Yiǧit et al. 2019). Then, randomly selected individuals were weighed on precision scales as 5 males and 5 females and then they were left to mate in a large cage. Those who did not mate within one hour (weight loss due to defecation) were left to feed, and a male and a female from the main population were randomly selected thereafter. The mean mating age for both sexes was 11 (min. 8-max. 15) days in P. inflatus and 12 days (min. 7-max. 23) in *P. sureyanus*.

In approximately 2 weeks, 24 matings from *P. inflatus* and 27 matings from *P. sureyanus* were achieved. The body weight of the males was quickly weighed immediately after mating. Then, the females were weighed with and without spermatophore. Non-virgin males and females were left to feed in separate cages for immunity studies for the next day. After the spermatophore was carefully weighed, the spermatophylax and ampulla were gently separated and weighed separately. All weights were measured to the nearest 0.1 mg. Afterwards, the ampulla were broken in 1 ml of water prepared in advance and homogenised with the help of an injector, and then counted with a hemocytometer, with 5 repetitions for each individual. Before each count, ejaculate and its contents were homogenised with an injector.

Immune parameters were determined by measuring Phenoloxidase (PO) and antibacterial (lysozyme-like activity, LY) activities, which play an important role in many immune responses. In the innate immune system of insects, PO is found in active and inactive forms in the hemolymph, and a humoral immune response is demonstrated against parasitoid and pathogens with cytotoxic products generated by the melanization pathway (Cerenius and Söderhäll 2004). LY, another important humoral immune response, is particularly effective against gram-positive bacteria (Adamo 2004). For immune assays, one day after successful mating, the insects were anaesthetised with CO2, 5μl of haemolymph were removed from abdominal sternites (via lateral pleural line between II. and III. abdominal sternite) using a Hamilton microsyringe (10μl, Model 701 RN SYR), 30-gauge, Small Hub RN NDL). It was tested whether the amount of withdrawn hemolymph was sufficient to measure PO and LY activities (Sevgili 2016). The sample was centrifuged with phosphate buffer saline and directly frozen at -20°C for PO and LY assays. To acquire the estimates of PO and LY, activities were assessed following the protocols of related literatures (Fedorka et al. 2013; Fedorka and Sevgili 2014; Sevgili, 2019). With a microplate reader (Multiscan FC, Thermo Fisher Scientific, at 25ºC), 10 measurements were taken at five-minute intervals using 495 nm for PO and 450 nm for LY. To avoid confusion in the analysis of LY, which normally has negative values, ΔOD values were calculated as positive absolute values (Fedorka and Sevgili 2014).

### Statistics

Statistical analyses were conducted using RStudio (RStudio Team 2021). Normality tests of the data were done with Shapiro-Wilk’s test. Both Phenoloxidase (PO) and Lytic Activities (LY) were not normally distributed, and they could not be transformed. Body and ampulla weight of the species were not distributed normally. Hence, non-parametric tests were used to compare immune and some mating parameters of both species and their sexes. Figures were generated in “*ggplot2*” (Wickham 2009). To understand the factors that may affect the spermatophore characteristics a linear mixed effect model (“*nlme*” package in R) was used (Pinheiro et al. 2021). In this statistic, individuals were considered as random factors, while body size (weight and hind femur length), development duration (the number of time or the number of days between individuals hatch from eggs to their adult stage), species and age were considered as fixed factors.

## Results

### Morphogenesis and body size

Under the same development conditions, *P. sureyanus* individuals completed their developmental process earlier than *P. inflatus* and reached adult stage (Fig. 2). There were differences in the duration of morphogenesis between males and females of the species, and males entered the adult stage earlier than females in both species (Fig. 2). There was no significant difference between male body weights and posterior femur lengths, which is another important indicator of body size, in both species (Fig. 3A-B). Males of *P. sureyanus* produced larger ampullae and greater numbers of spermatozoa, whereas spermatophore weights did not differ for the two species (Fig. 3C-E). Males in both species transferred similar rates of spermatophore, while P. *inflatus* invested proportionally larger spermatophylax (Fig. 4A-B). In contrast, *P. inflatus* transfered an ampulla containing a much smaller ejaculate relative to body weight (Fig. 4C). In P. inflatus, the spermatophylax surrounding the ampulla (right after mating, the female consumes first the spermatophylax then the ampulla) was found to be proportionally larger (Fig. 4D).

**Fig. 2.**
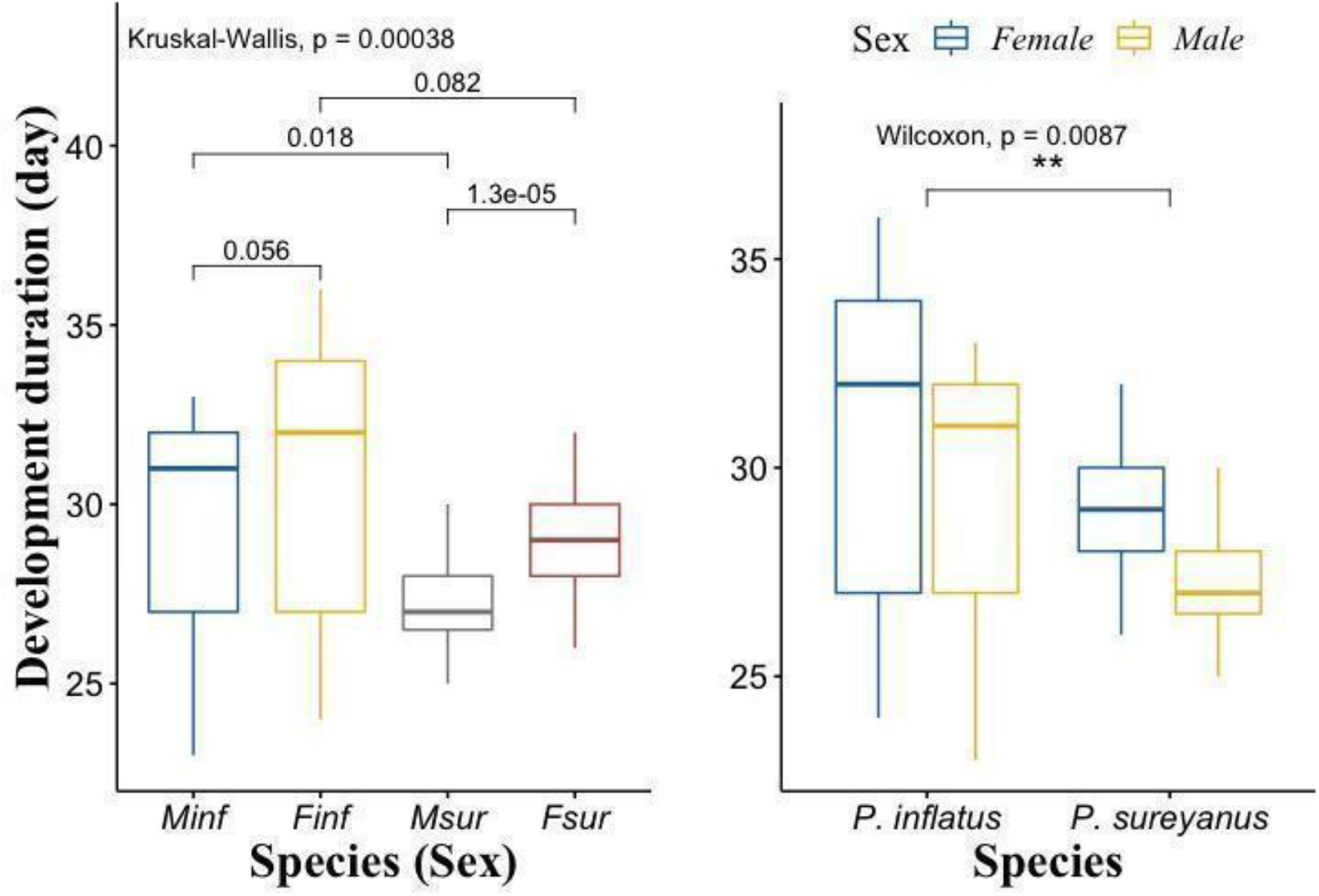
The development time was different in *P. inflatus* and *P. sureyanus* species. Although males reached earlier adult period than females in *P. inflatus*, no statistical difference was found. However, in *P. sureyanus*, males clearly matured much earlier than females. While there was no difference between females of the two species in terms of development time, the statistical difference between males was remarkable (Minf: *P. inflatus*-male; Finf: *P. inflatus*-female; Msur: *P. sureyanus*-male; Fsur: *P. sureyanus*-female).

**Figure 3.**
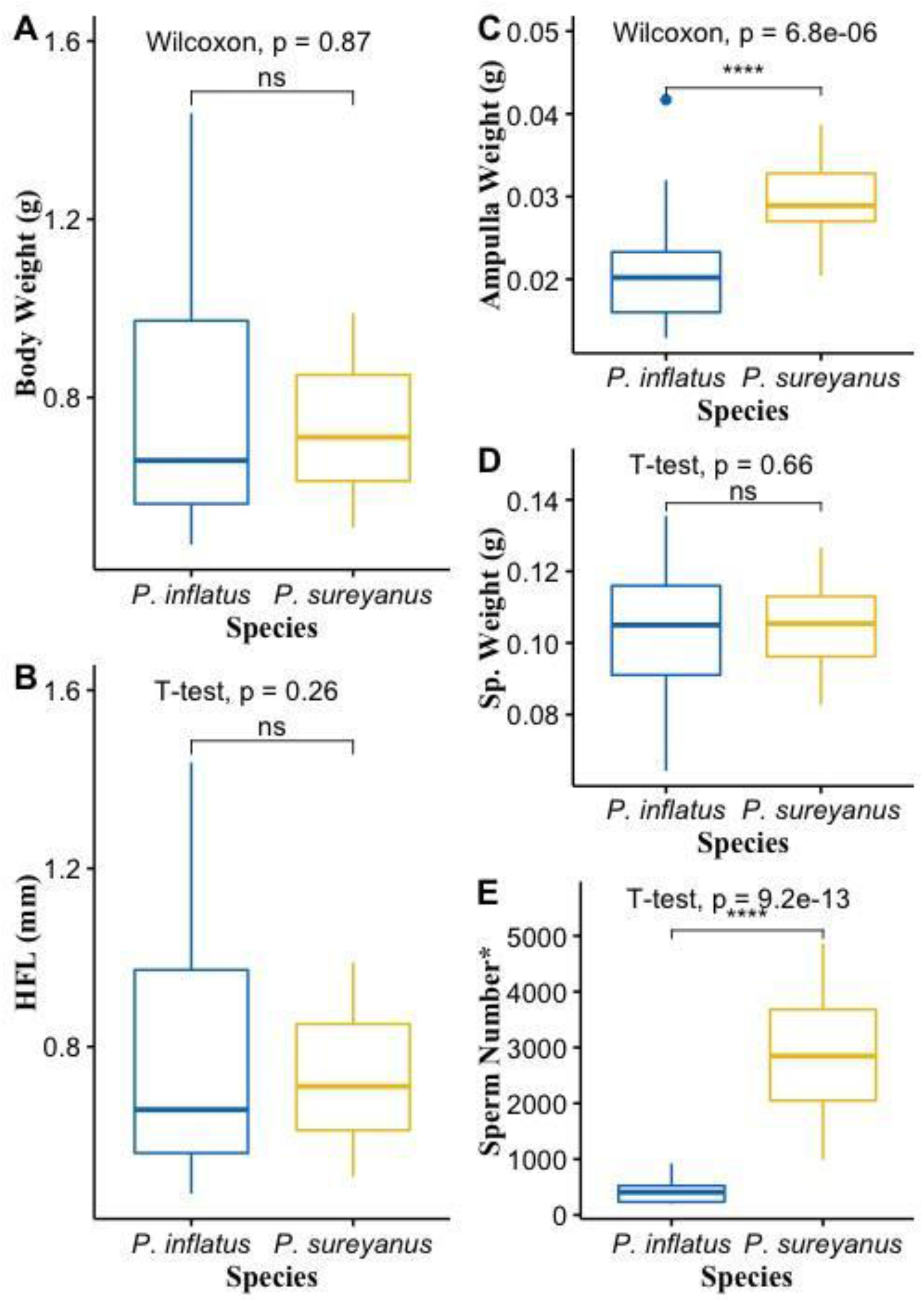
Different tests were used for normal and non-normally distributed parameters. *P. inflatus and P. sureyanus* males were compared for body size and some spermatophore parameters.

**Figure 4.**
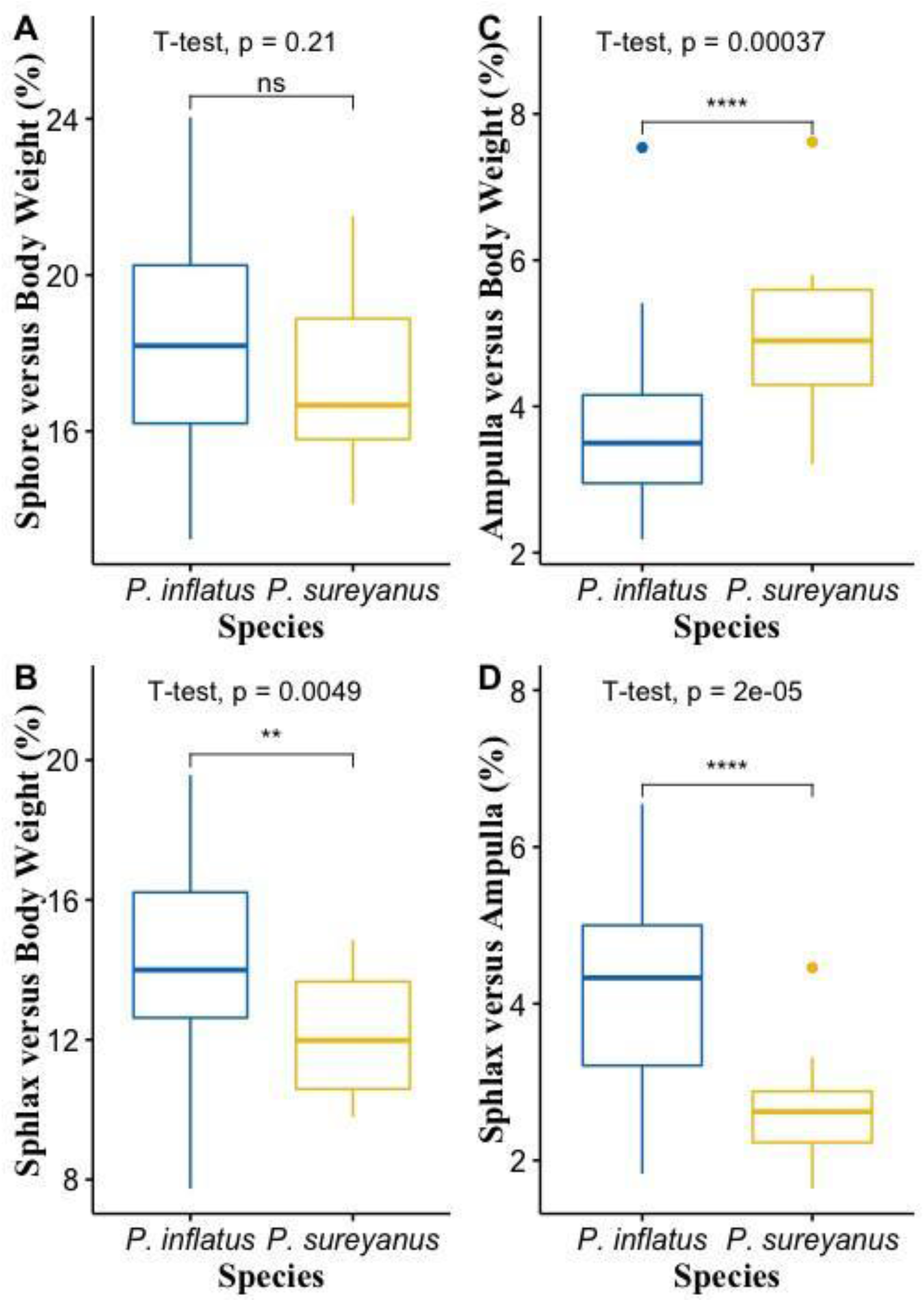
Proportional comparison of some spermatophore data in males of *P. inflatus* and *P. sureyanus*. The differentiation in spermatophylax production between these two species is particularly remarkable.

### Spermatophore

While spermatophore and spermatophylax weights were affected by the body size of the male for both species, it was determined that the ampulla containing the sperm was independent of body size (Table 1). It was observed that the difference in duration of morphogenesis between species had no effect on spermatophore weight and absolute sperm count (Table 1). Ampulla and sperm allocation were found to be a strong signal that the species might be affected by mate searching strategies (Table 1).

**Table 1.**
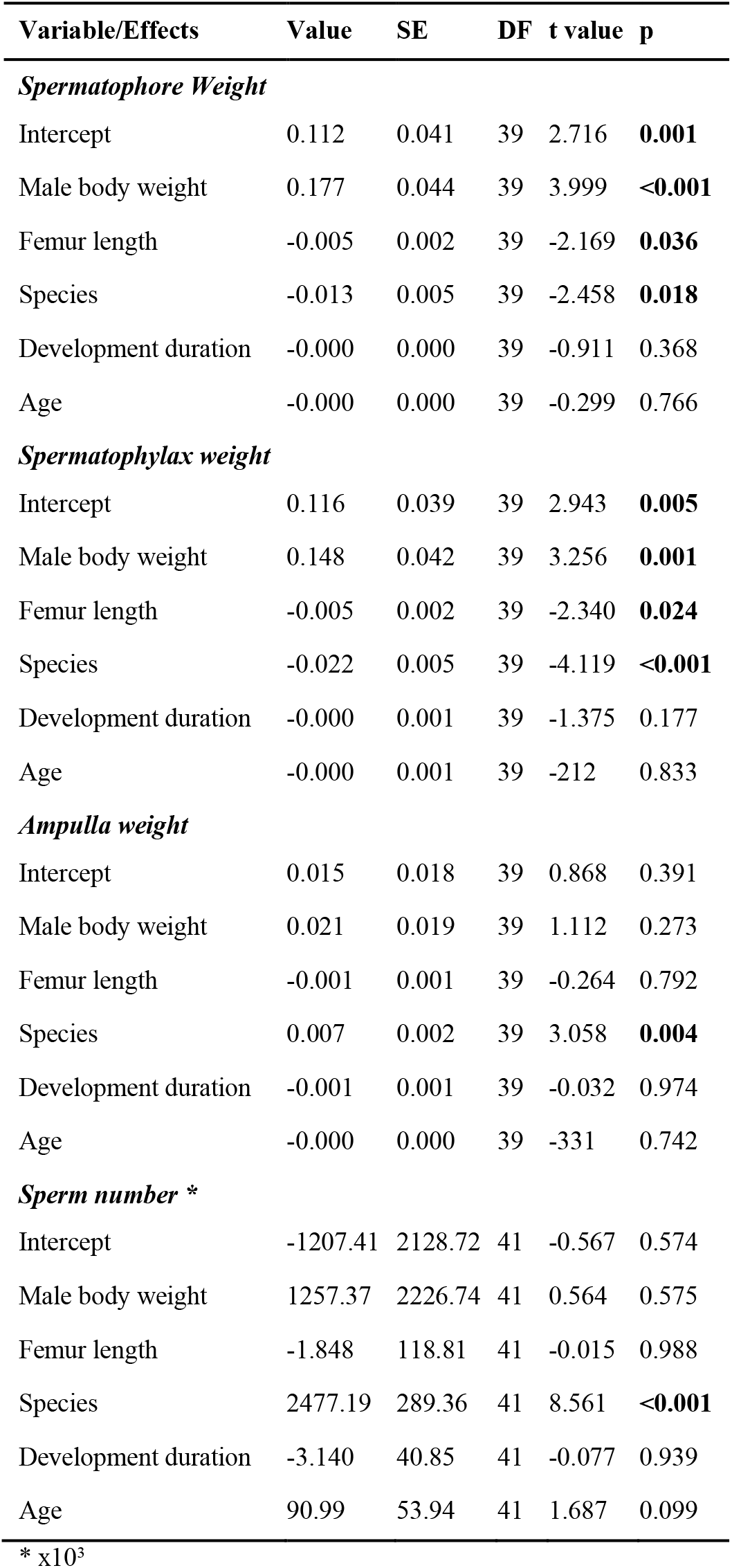
Results of Linear mixed-effects model fit by REML (Library “*nlme*” (Pinheiro et al. 2021), Male ID as random factor) testing the effects of male body weights, length of right hind femur, species, development duration and age on spermatophore characteristics.

| Variable/Effects | Value | SE | DF | t value | p |
| --- | --- | --- | --- | --- | --- |
| ***Spermatophore Weight*** |  |  |  |  |  |
| Intercept | 0.112 | 0.041 | 39 | 2.716 | **0.001** |
| Male body weight | 0.177 | 0.044 | 39 | 3.999 | **<0.001** |
| Femur length | -0.005 | 0.002 | 39 | -2.169 | **0.036** |
| Species | -0.013 | 0.005 | 39 | -2.458 | **0.018** |
| Development duration | -0.000 | 0.000 | 39 | -0.911 | 0.368 |
| Age | -0.000 | 0.000 | 39 | -0.299 | 0.766 |
| ***Spermatophylax weight*** |  |  |  |  |  |
| Intercept | 0.116 | 0.039 | 39 | 2.943 | **0.005** |
| Male body weight | 0.148 | 0.042 | 39 | 3.256 | **0.001** |
| Femur length | -0.005 | 0.002 | 39 | -2.340 | **0.024** |
| Species | -0.022 | 0.005 | 39 | -4.119 | **<0.001** |
| Development duration | -0.000 | 0.001 | 39 | -1.375 | 0.177 |
| Age | -0.000 | 0.001 | 39 | -212 | 0.833 |
| ***Ampulla weight*** |  |  |  |  |  |
| Intercept | 0.015 | 0.018 | 39 | 0.868 | 0.391 |
| Male body weight | 0.021 | 0.019 | 39 | 1.112 | 0.273 |
| Femur length | -0.001 | 0.001 | 39 | -0.264 | 0.792 |
| Species | 0.007 | 0.002 | 39 | 3.058 | **0.004** |
| Development duration | -0.001 | 0.001 | 39 | -0.032 | 0.974 |
| Age | -0.000 | 0.000 | 39 | -331 | 0.742 |
| ***Sperm number \**** |  |  |  |  |  |
| Intercept | -1207.41 | 2128.72 | 41 | -0.567 | 0.574 |
| Male body weight | 1257.37 | 2226.74 | 41 | 0.564 | 0.575 |
| Femur length | -1.848 | 118.81 | 41 | -0.015 | 0.988 |
| Species | 2477.19 | 289.36 | 41 | 8.561 | **<0.001** |
| Development duration | -3.140 | 40.85 | 41 | -0.077 | 0.939 |
| Age | 90.99 | 53.94 | 41 | 1.687 | 0.099 |

\* $\times 10^3$

### Immunity

PO and LY activities differed between sexes and within sexes for both species (*P. inflatus*, n= Male 23, Female 23; *P. sureyanus*, n= Male 27, Female 27) (Table 2). There were no significant relationships between PO and LY in males for both species (Spearman corr., *P. inflatus*, R= 0.29, p= 0.49; *P. sureyanus*, R= 0.21, p= 0.57). When males and females of both species were included in the analysis, only PO activity differed between the two species, while no significant difference was found in LY activities (Wilcoxon, PO, p<0.001; LY, p= 0.22). When evaluated in terms of sexes within the species, LY did not differ between sexes in either species (Wilcoxon, p= 0.44-*P. inflatus*; p= 0.51-*P. sureyanus*). However, PO did not differ between species in males, while LY was significantly weaker in *P. sureanus* (Figure 5). When both species were evaluated amidst themselves, PO activity was weaker in females in *P. inflatus*, but stronger in females of *P. sureyanus* (Wilcoxon, p<0.001; *P. inflatus* p= 0.003, *P. sureyanus* p= 0.012).

**Table 2.**
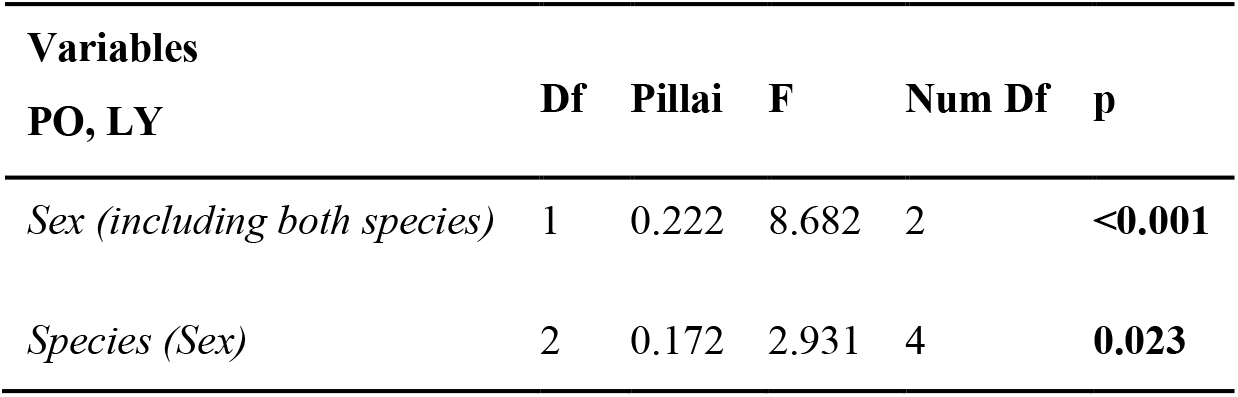
In *P. sureyanus* and *P. inflatus*, the sexes and between the sexes of the species were compared with PO and LY MANOVA ((cbind(PO and LY) in R)).

**Figure 5.**
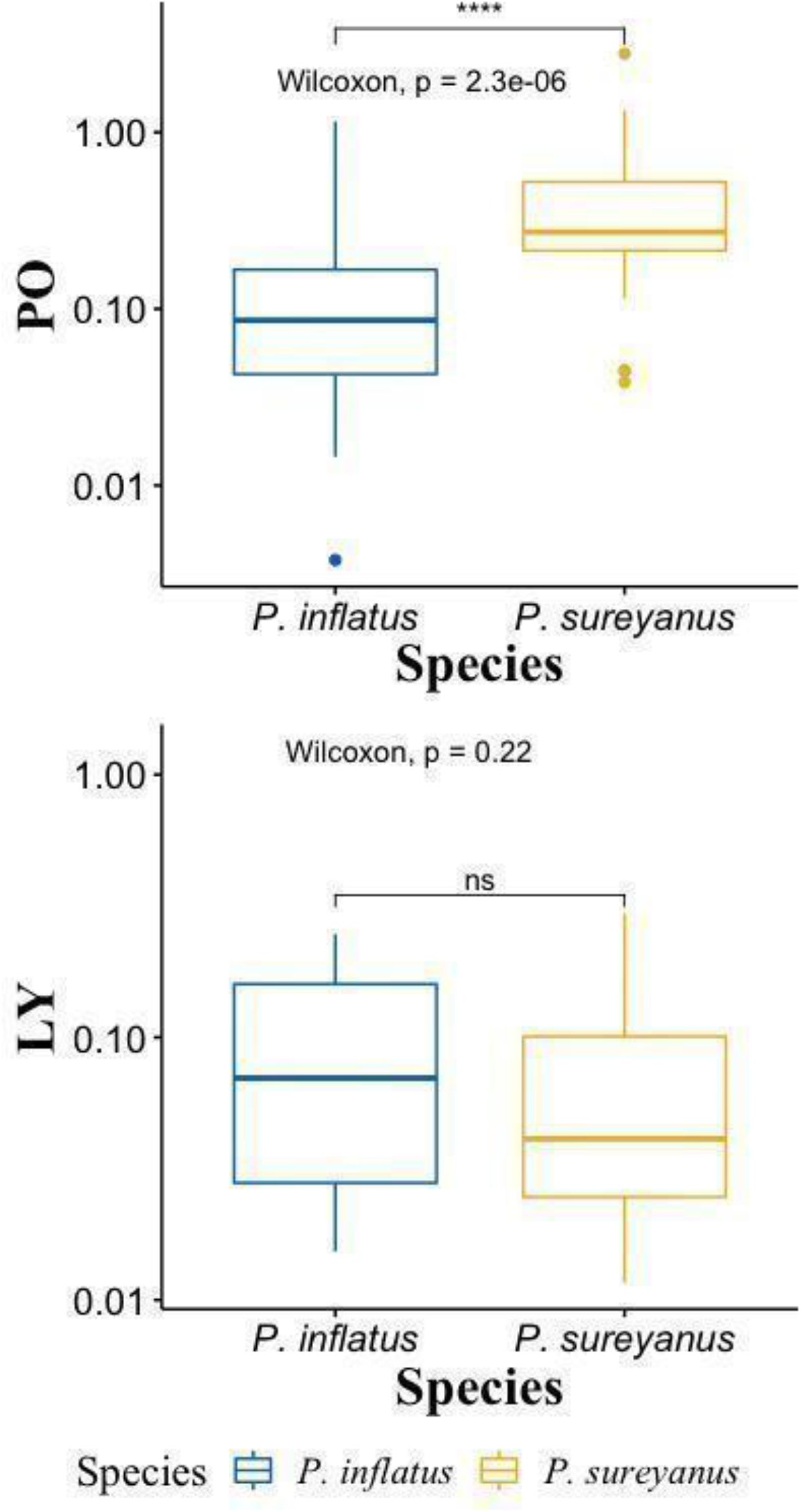
Box-plot visualisation of PO and LY activities for both species including all sexes.

**Figure 6.**
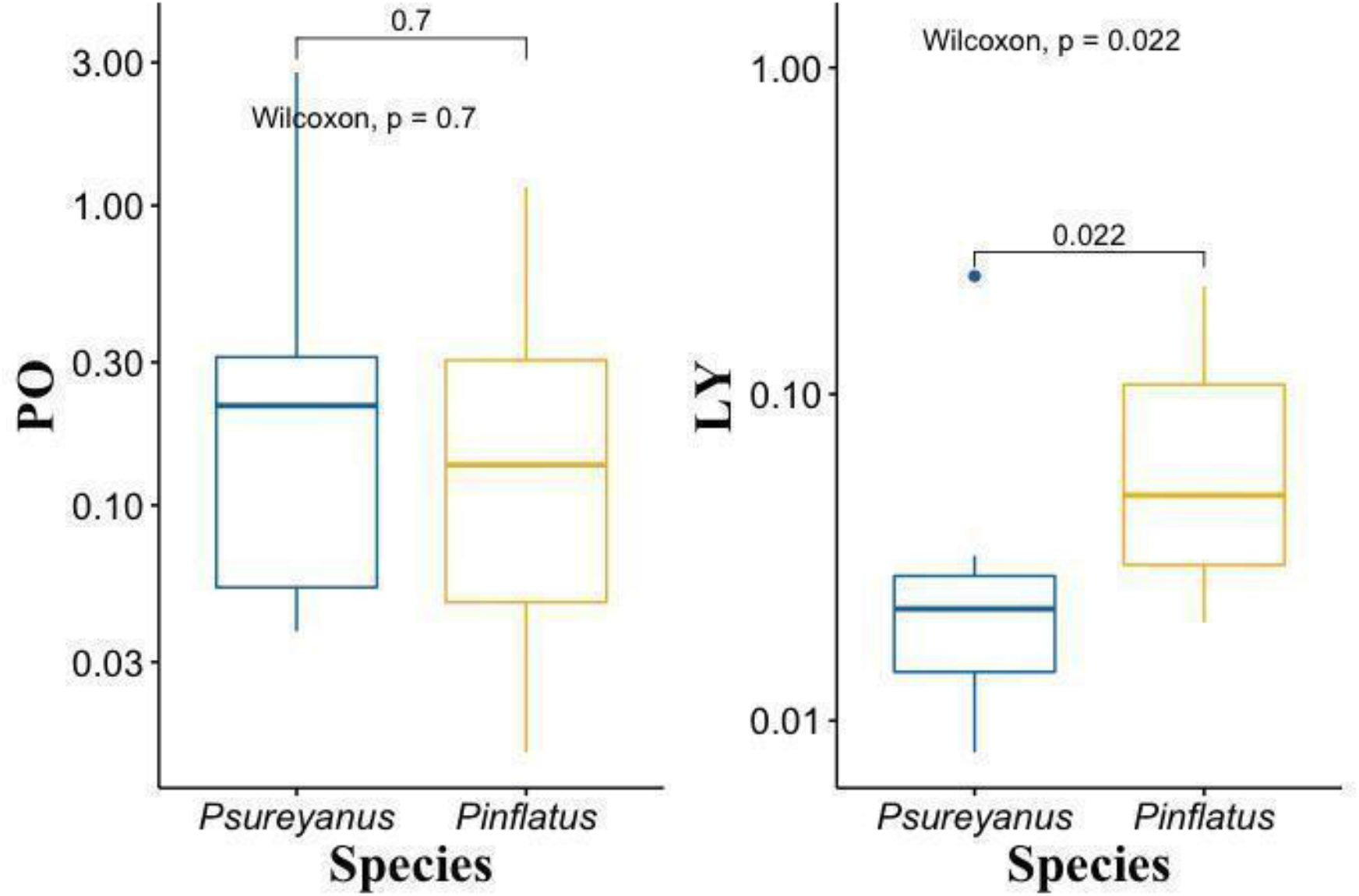
TPO and LY activities were only considered for the males, PO does not differ between species, while LY is markedly different40

## Discussion

### Morphogenesis and body size

It is known that the relationship between morphogenesis and body size in Orthoptera is determined by many environmental (eg temperature, precipitation, humidity, altitude, nutrition) and genetic factors (Berner and Blanckenhorn 2006; Whitman 2008; Çaǧlar et al. 2014). For example, it is known that bush-populations at high altitudes have a shorter developmental period and a smaller body than those living in low areas (Orthoptera Whitman 2008; Poecilimon veluchianus Eweleit and Reinhold 2014). Both species in this study developed under the same environmental conditions and their nymphal stage numbers (5) were the same. It has been revealed that there is a positive relationship between longer development period and body size in some insect species (Sõber et al. 2019).

In this study, although the duration of morphogenesis was different in both species, the fact that body sizes (posterior femur length) were not different indicates that body sizes are mostly determined by genetic factors. However, it is clear that a more detailed study is needed to reach an acceptable conclusion on this issue. In both species, it is a common and well-known phenomenon in insects that males complete their development earlier than females (Jarosik and Honek 2007). While males reached the adult stage much earlier in *P. sureyanus*, there was no statistically significant difference between the sexes in *P. inflatus*. Since females are already mute in *P. sureyanus*, only males begin to call a song a few days after eclosion. This is typical for males as no sound signal response is expected from females ready to mate. On the other hand, since the *P. inflatus* male expects a response from the female, it can be said that the coincidence of the two sexes reaching the adult stage is more significant.

### Spermatophore

In this study, the spermatophore characteristics and some immune activities of two species with differences in mate searching strategies were compared, and no significant difference was found in spermatophore investment, unlike the theory proposed by McCartney et al (2012). However, in accordance with the theory, in the system where the female seeks the male (*P. sureyanus*), the male produces larger ampullae and more sperm. In the system where the male seeks the female, *P. inflatus* produces larger spermatophylax compared to the male body weight. Also, in this species, the larger ampulla was surrounded by larger spermatophylax. This finding supports the ejaculate protection hypothesis(McCartney et al. 2008; Lehmann 2012).

The results of this study indicate that the strategy in spermatophylax investment is related to the size of the ejaculate rather than the sex-searching system. On the other hand, the fact that the ampulla is significantly larger in terms of body weight ratio in the system where the female seeks for the male supports the existing theory. Perhaps in this system, ejaculate investment and the selection of spermatophylax to protect the ejaculate are more decisive than its weight - which is considered the main determining factor in the spermatophore transfer strategy in this system. Because spermatophylax production is the costliest part of the spermatophore (Heller et al. 1998; McCartney et al. 2012), it has been reported that amino acids such as glycine in the content of the spermatophore consumed by the female suppress the female’s desire to mate (Gordon et al. 2012) (Gordon et al., 2012), as well as limit the mating frequency of the male (Reinhold and Helversen 1997; Uma and Sevgili 2014).

It is known that in bush-crickets, the male needs a few days to produce fresh spermatophores and the sperm number increases in subsequent matings depending on the age, but after about three weeks, the sperm increase stops (Uma and Sevgili 2014). The male’s age, which is an important factor in the data discussed in the theory, was not included in the model. While there is some evidence that the male performs spermatophore transfer according to the female’s body weight in bush-crickets, there is a lack of studies on whether the body size of the female with whom the male mates is effective in adjusting the spermatophore size and sperm number in the two different mate searching strategies in the genus *Poecilimon*. In *P. jonicus jonicus*, a species in which the female responds acoustically to the male, there is no evidence to support that the male adjusts its sperm number according to the female’s weight (Sevgili and Reinhold 2007). Therefore, it will be inevitable for the theory to undergo some changes after studying the species in detail. In populations, many factors influence the behaviour of searching and finding the sex ready to mate. Although many models have been proposed as to which sex is more dispersed in the population, there are many difficulties to fully explore and explain the unknown aspects of the system (Shaw and Kokko 2014).

After mating, females are presumed to be more stable than males because they feed on spermatophores for a long time. However, depending on the population and the density of the predators, the mate searching strategy requires that the sex-biased activity in the population be variable. In particular, the male’s calling song signals are beacons for predators (e. g. *Therobia leonidei*, Diptera), and males are more susceptible to phonotactic predation than females (Lehmann and Heller 1998). On the other hand, after a costly spermatophore transfer, it takes a few days for males to be ready to mate again, which is thought to be more stable during this time period (Vahed 1998; Kerr et al. 2010; Uma and Sevgili 2014).

The differences in ampulla weight, sperm number, and immune activities between sexes and species can be identified as an evolutionary consequence of differences in mate searching strategies. It has been reported that mobility and survival rates differ between sexes in *Poecilimon veluchianus* and *P. affinis* species, which show different searching roles (Heller 1992b). However, it should be taken into account that the habitat and vegetation differences in which the species live may also affect the costs arising from their mate searching roles. On the other hand, factors such as multiple mating being common due to sperm competition and the role of nuptial gift to females in reproductive success (Gwynne 1988) may have been effective in the divergence in the mate-searching role in bush-crickets. Partially different from the hypothesis proposed by McCartney et al (2012), spermatophore produced by *P. inflatus*, which fit the model in which the male seeks the female, did not differ from *P. sureyanus*, while a significant difference was found in the ampulla weight and sperm number. This finding suggests that it may be controversial whether the theory proposed by McCartney et al (2012) is valid for the entire content of the spermatophore. The obvious finding is that, according to what the results indicate males may tend to produce less sperm in the system where the male seeks the female. There is a wide variation in spermatophore and ampulla weight and sperm number in *Poecilimon* species (McCartney et al. 2008; Sevgili, 2016). It should be noted that the difference in sperm transfer seen in these two species is shaped not only by the sex roles in mating, but also by both proximate and ultimate factors.

### The Immunity

In animal species (from insects to mammals), immune responses are highly diversified between sexes, and in general, both innate and adaptive immunity are typically stronger in females than in males (Rolff 2002). The immunity of sound signallers may be weaker, a sex (usually males) allocates some of its energy resources on reproductive efforts such as advertisement signals to other sex (bioacoustics, pheromones) and visual exaggeration of color patterns (mostly in diurnal butterflies). For example, in *Heliothis virescens* (Lepidoptera), males are stronger in terms of immunity as females are the sexual signallers (Barthel et al. 2015). There is a trade-off between mating behaviours and the costs necessary for immunity. For example, males (*Drosophila melanogaster*) exposed to larger numbers of females were less effective against bacterial defence (McKean and Nunney 2001). It has been reported that males producing larger spermatophores in *Gryllodes sigillatus* have weaker immunity (Kerr et al. 2010). The difference in spermatophore and some immune activities in the two *Poecilimon* species, which were the subject of this study, whose activities in reporting and determining the locations of potential mates were asymmetrical, also provided evidence in favour of this trade-off. When resources are limited, it is possible for individuals to invest more in, therefore pronounce one feature while partly renouncing others. For example, nutrition restrictions negatively affect immunity and sperm allocation (Simmons 2012).

A trade-off between spermatophore investment and immunity parameters has been demonstrated in *G. sigillatus*, where males producing larger spermatophores exhibit lower immunity (Kerr et al. 2010). In *Ephippiger diurnus* (Orthoptera: Bradyporinae), which conforms with the system in which the males actively seek females, a negative correlation was found between the spermatophore size and the number of call syllables and encapsulation, indicating a trade-off between reproductive and survival traits (Barbosa et al. 2016). The results of this study revealed that PO activity was more plastic than LY activity and was affected by the sex role reversal in mate-searching, but LY activity did not differ significantly between sexes in either species. In general, females have a stronger immunity than males (Steiger et al. 2011; Kelly et al. 2018; Sevgili 2019b). It is an interesting finding that the immune activity is more in favour of the male in the system where the females actively seek the males, than the one in which the male is the seeker. Thus, it is possible to say that there may be trade-offs between the energy spent for spermatophore production and the energy spent for mate-searching. Survival traits such as immunity may have been traded off not only with spermatophore production but also with the effort to find a mate. It has been reported that, in a cricket species, males who spend more energy for immunity take longer to mate and spend less time producing sound signals (Sagebrush crickets Leman et al. 2009).

As a result, it has been shown that a difference in mate searching based on whether the female responds to male calling songs bioacoustically for reproduction, may play a decisive role on both spermatophore and immune activities. The results of the study also indicate that there may be trade-offs between the costs related to survival traits and reproduction. It has been shown that there may be a trade-off between immune activities and differences between sexes. On the other hand, the data suggest that the duration of morphogenesis does not have an effective role on reproduction and survival traits in holometabola bush-crickets. The theory put forward by McCartney et al (2012) was partially supported for *P. sureyanus* (female search) and *P. inflatus* (male search).

## Acknowledgement

I would like to thank Ebru Kıran Özdemir for helping with the maintenance of the bush-crickets.

